# Aquatic insects of a lowland rainforest in Papua New Guinea: assemblage structure in relation to habitat type

**DOI:** 10.1101/028423

**Authors:** Jan Klecka

## Abstract

Papua New Guinea is one of the most valuable tropical regions but ecological research of its freshwater invertebrates has been lacking. The goal of this paper is to evaluate the species richness, diversity and structure of aquatic insect assemblages in different habitats in the Wanang River catchment in a well-preserved lowland rainforest. Assemblage structure was studied on two spatial scales – in different habitats (river, streams and stagnant pools) and in three mesohabitats in the river (slow and fast sections and submerged wood). The results show that headwater streams had the highest morphospecies diversity, while the river had the highest insect abundance. Slow and fast sections of the river differed both in terms of insect abundance and diversity. Furthermore, a number of unique wood-associated species was found on submerged wood. The most notable feature of the assemblage structure was scarcity of shredders and dominance of predators. However, predatory beetles, bugs and dragonfly larvae exhibited contrasting habitat preferences. This study shows that Papua New Guinean lowland rainforests host diverse and distinctly structured freshwater insect assemblages.

## Introduction

Better insight into the structure and functioning of freshwater communities of so far insufficiently known regions is needed to test the generality of empirical patterns and theories based on intense research in temperate regions. Our knowledge of tropical freshwater ecosystems has progressed significantly in the last decades but has been restricted only to a handful of geographical areas (Boyero et al. 2009). Tropical waters may differ form their temperate counterparts in the diversity and composition of insect communities with potentially important implications for ecosystem functioning. Most groups of aquatic invertebrates seem to have higher species richness in tropical than in temperate regions (Abel 2008; Balian et al. 2008a, b; Pearson & Boyero 2009). Nevertheless, some taxa, e.g. Ephemeroptera, Plecoptera and Trichoptera, deviate form this pattern, although the paucity of tropical data precludes drawing definitive conclusions (Vinson & Hawkins 2003; Barber-James et al. 2008; de Moor & Ivanov 2008; Fochetti & Tierno de Figueroa 2008; Pearson & Boyero 2009). Low diversity of stoneflies and caddisflies in the tropics is responsible for the overall scarcity of shredders (e.g. Dudgeon 1994; Yule 1996c; Boyero et al. 2009; Li & Dudgeon 2009), which may substantially decrease the rate of leaf litter processing in tropical waters and alter the paths of energy flow in their food webs (Graça 2001; Boyero et al. 2009).

Environmental characteristics affect the abundance and species composition of aquatic insects at the habitat scale (stagnant vs. running waters) as well as at the mesohabitat scale (e.g., riffles and pools in rivers). Aquatic insects in temperate rivers are usually more abundant and have higher total biomass and species richness in riffles compared to pools (e.g. Slobodchikoff & Parrot 1977; Brown & Brussock 1991; Lemly & Hilderbrand 2000; Cheshire et al. 2005), although some studies found the opposite pattern (McCulloch 1986). The variation of the taxonomical composition of aquatic insect assemblages between these two mesohabitats is coupled with the variation of the relative abundance of functional feeding groups. The availability of food is probably responsible for such a variation; e.g., scrapers are abundant in riffles with stones overgrown by periphyton, while shredders are associated with thick layers of leaf litter accumulated at the bottom of slow sections and with leaf packs retained among stones in riffles (e.g. Angradi 1996). A special type of mesohabitat common especially in rivers is submerged wood. Its presence affects invertebrates indirectly by increasing habitat complexity and creating accumulations of detritus (Lemly & Hilderbrand 2000). Invertebrates associated with wood are little studied, but they can be fairly diverse, at least in some regions (Johnson et al. 2003). Testing the generality of these patterns in freshwater invertebrate community structure requires more data on tropical rivers and streams.

Papua New Guinea, a biodiversity hotspot and a prominent region in the research of tropical rainforests (e.g., Novotny et al. 2002), is surprisingly almost a blank sheet in freshwater ecology. So far, only Dudgeon (1990, 1994) studied the composition of aquatic insect assemblages in relation to riparian vegetation in several lowland streams. Yule (1995, 1996a, 1996b, 1996c) conducted detailed research on macroinvertebrates in fast flowing mountain streams in Bougainville Island. This small volcanic island is located ca. 600 km from the coast of Papua New Guinea and its geomorphology and habitats studied by Yule (1995, 1996a, 1996b, 1996c) are indeed very different from that of Papua New Guinean lowland rainforests.

I conducted a short-term field study to advance our understanding of the structure of freshwater assemblages in lowland rainforests in Papua New Guinea. My goal was to compare the species richness, abundance and taxonomical and functional composition of insect assemblages among major habitats available in the rainforest during the dry season. I analysed the differences among different habitats (river, streams and stagnant pools) and among mesohabitats in the river (slow and fast sections and submerged wood).

## Methods

### Study site

The fieldwork was carried out in the end of July 2008 (middle of the dry season) in the vicinity of Wanang Village in Madang Province, Papua New Guinea (05°13’85’’ S, 145°10’91’’ E; ca. 100 m a.s.l.; Fig. 1). The climate of the region is characterised by nearly constant high temperature and alternating dry and wet season once a year with a mean annual rainfall of 3107 mm (Supplementary Fig. S1). The study site was located in a large complex of lowland rainforests impacted only by small-scale agricultural activities of traditional human societies. Although the precipitation levels drop significantly in the dry season, some rainfall still occurs (Supplementary Fig. S1) and only some streams and pools dry out. The survey focused on the Wanang River (5 sites), small headwater streams (7 sites) and small pools (6 pools at 2 sites) in a narrow forested valley along ca. 5 km long stretch of the river (Fig. 1); detailed characteristics of individual sites are described in Supplementary Table S1.

**Fig. 1.**
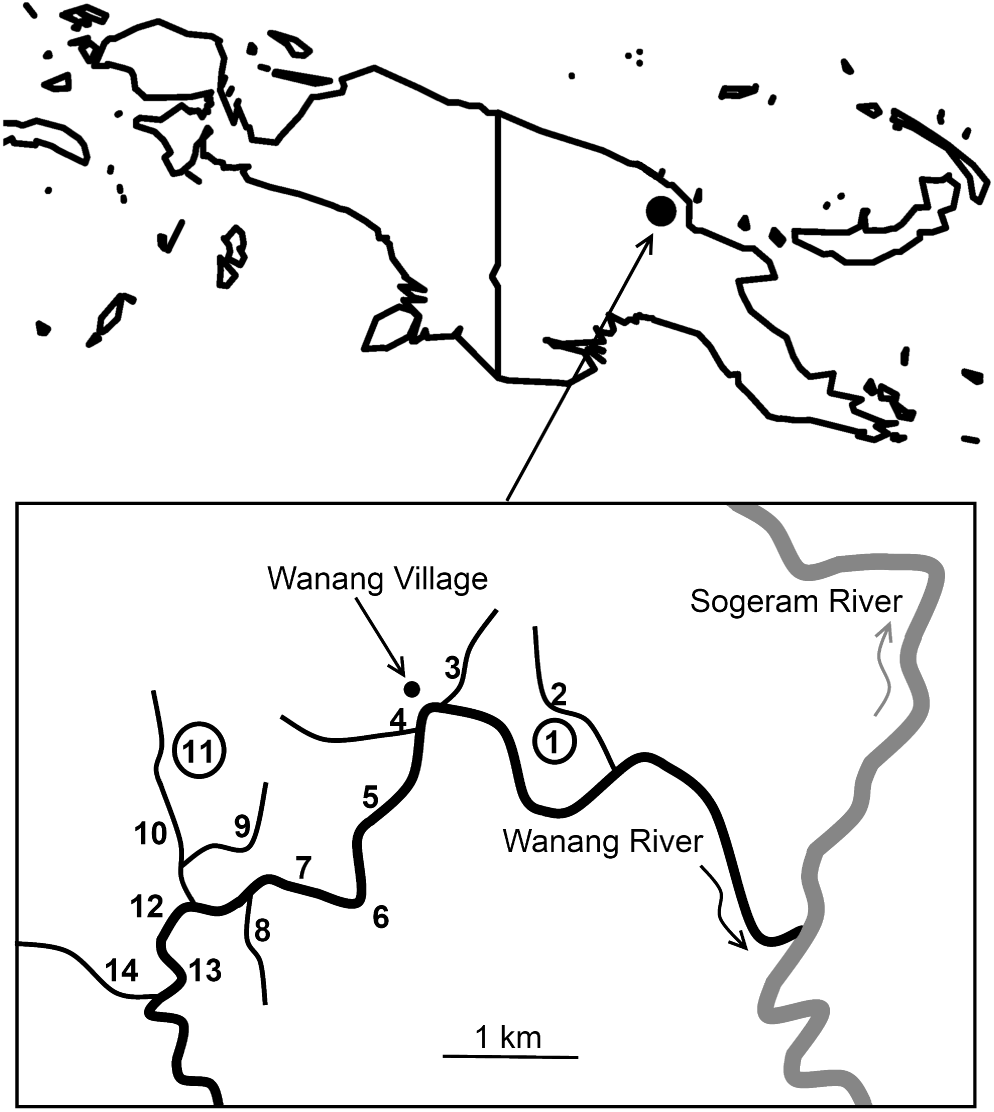
Map of the island of New Guinea with the position of Wanang Village and a detailed map of the study area with all study sites. The positions of the sites are marked by numbers that refer to site descriptions in Supplementary Table S1. The two numbers in circles (1 and 11) are stagnant pools; the remaining sites are the Wanang River (5, 6, 7, 12 and 13) or streams (2, 3, 4, 8, 9, 10 and 14).

Individual sampling sites are listed in Supplementary Table S1 and their spatial arrangement is depicted in Fig. 1. Shallow riffles (depth <20 cm, current speed<0.5 ms^−1^) and deep slow sections (depth <1 m) alternated in the Wannang River. Three mesohabitats were sampled at all sites in the river – slow sections, fast sections and submerged wood (fragments of trunks and branches of trees) deposited in slow sections. River mesohabitats are hereafter referred to as slow and fast sections because not only running waters but also stagnant waters were sampled; pool is used only for a small stagnant water body to avoid confusion. Apart from the river, seven streams were sampled; all of them were tributaries of the Wanang River (Fig. 1). The streams had steeper slope than the river but the current speed was similar to that of the river’s fast sections. All stagnant pools (6 pools at 2 sites) found in the vicinity of the focal stretch of the Wanang River were sampled (excluding several polluted pools in the Wanang Village).

### Sampling and processing material

Three samples were obtained at each site in the river, one for each of the three different mesohabitats (fast and slow section and submerged wood). No mesohabitat division was made in streams and pools because they were too small to allow us to differentiate different mesohabitats; wood was not sampled because it was rare. We used a semi-quantitative sampling protocol consisting of 60 minutes of sampling (20 minutes by three people or 15 minutes by four people). We disturbed the bottom and collected insects using steel strainers (diameter 20 cm, mesh size 0.75 mm). The strainers were used mainly because it would not have been possible to use larger benthic handnet or Surber sampler in small streams and shallow parts of the river’s fast sections. Submerged wood was sampled by removing tree branches and fragments of trunks out of water and manually collecting insects from the surface of the wood over the same time period as above. In total, 24 samples were collected (15 in the river – 5 sites × 3 mesohabitats, 7 in headwater streams and 2 in pools).

All collected insects were stored in 70% ethanol and were identified and enumerated in the lab; samples were not subsampled. Most specimens were identified to families and morphospecies (mostly according to Williams (1980) and Yule & Sen (2004)), which is a standard approach used by entomologists working in insufficiently known tropical regions. Higher precision of specimen identification was not possible because of the lack of taxonomical knowledge of Papua New Guinean aquatic insects. Larvae and adults of one dominant species of a beetle from family Elmidae were associated based on co-occurrence in samples and corresponding body size. In all other cases, larvae were sorted into morphospecies independently of adults. Larvae of Heteroptera could be easily associated with adults in all cases. All morphospecies were classified to functional feeding groups based mainly on family level information on feeding habits in the literature (mostly according to Williams (1980) and Yule & Sen (2004)), which is crude but widely used approach (e.g. Angradi 1996, Boyero et al. 2009, Flores et al. 2011). Species collecting fine particulate organic matter were included in the group of collectors because further division in filtrators and gatherers would be unreliable. The presence of beetles, mostly of the family Elmidae, associated with wood requested including a special group, which is missing in most comparable studies (but see Johnson 2003). These beetles feed on decaying wood or associated fungi and bacteria (e.g., Williams 1980, Moog 2002, Johnson 2003); they are called here xylobionts.

### Data analyses

Two levels of data analysis were considered: an analysis of differences among habitats (river, streams and pools) and an analysis of differences among mesohabitats in the river (fast and slow sections and submerged wood). In the first of these analyses, the samples from different sites were regarded as replicates for one of the three habitats within the sampling area. To avoid pseudoreplication and enable comparison among habitats, river samples from the fast and slow sections at individual sites were pooled. Wood samples were excluded and used only in the analysis comparing mesohabitats in the river because wood was not sampled in the streams and pools.

The insect abundance, species richness and diversity were compared among habitats using generalized linear models (GLM) with quasi-poisson distribution for abundance and species richness and normal distribution for diversity and equitability indices in R 2.9.2 (R Core Development Team 2009). An analogous analysis was performed to compare abundance, species richness, and diversity indices between mesohabitats in the river. In these analyses, site was used as a random factor and generalized linear mixed effects models were used with normal distribution for diversity and equitability indices and Poisson distribution for abundance. The significance of mesohabitat (fixed factor in the analysis) was tested by a log-likelihood test comparing models with and without this predictor. These analyses were conducted using lme4 package for R (functions lmer and glmer; Bates et al. 2014a, 2014b). Rarefaction analysis was performed using an individual based rarefaction procedure in Analytic Rarefaction 1.3 (Holland 2003) to compare morphospecies richness among different habitats as well as among mesohabitats in the river because the number of individuals collected differed strongly, which precludes a direct comparison of morphospecies richness. Morphospecies diversity was evaluated using the Shannon’s diversity and equitability index and by the Simpson’s index. Shannon’s diversity (Shannon & Weaver 1949) is derived from the information theory and quantifies the entropy of a system. Shannon’s equitability can be calculated by dividing the Shannon’s diversity index by ln(*S*), where *S* is the number of species. The Simpson’s index (Simpson 1949) is equal to the probability that two randomly chosen individuals belong to the same species; the inverse of this probability ranges from 0 (minimal diversity) to the total number of species (maximal diversity) in a sample; this form of the index was used here. Both Shannon’s and Simpson’s diversity index are affected by both species richness and relative abundances, while Shannon’s equitability corrects for differences in species richness and represents only information on relative abundances of species (Magurran 1988).

To compare species composition of different habitats and mesohabitats, redundancy analysis (RDA) was used based on guidelines provided by Šmilauer & Lepš (2003). Standardization by sample norm was used to test for differences in species composition (i.e. relative abundance) between habitats rather than differences in absolute abundance. Species data were centred and standardized. This procedure, also called normalization, means that the abundances of each species are transformed to have mean equal to zero and unit variance. Consequently, all species have the same weight in the analysis (Šmilauer & Lepš, 2003). The analysis was computed in CANOCO 4.5 (ter Braak & Šmilauer 2002). Monte Carlo test with 9999 permutations was used to test significance in both cases. In the case of mesohabitats in the river, site identity was used as a covariable and permutations were carried out within sites. The similarity of samples within individual habitats and mesohabitats was estimated by the Chao-Sorensen index corrected for undersampling bias (Chao et al. 2005) in EstimateS 8.0.0 (Colwell 2006).

## Results

### Richness and diversity

In total, we have collected 78 morphospecies of aquatic insects in 24 samples (Supplementary Table S2). The average number of morphospecies per sample was 25 across different habitats (range from 16 to 41) and 14 across river mesohabitats (range from 7 to 28). The most diverse groups were Coleoptera (27 morphospecies), Hemiptera (16) and Odonata (12). Diptera were represented by 11 morphospecies but their abundance was surprisingly very low (Supplementary Table S2).

Mean insect abundance per sample was higher in the river (163 individuals, range from 125 to 205) than in the streams (91, range from 62 to 128) and pools (83, range from 77 to 89) (Table 1). The highest number of morphospecies per sample was also found in the river (32 morphospecies, range from 29 to 41). Rarefaction curves (Fig. 2) allow estimating total species richness in individual habitats. The total number of species is much lower in stagnant pools (27) than streams (59) and rivers (59). However, the number of individuals collected in streams was more than three times lower than in rivers and the rarefaction curve for streams does not reach an asymptote (Fig. 2), so there is a potential for collecting more species in streams unlike in the river and pools. At the level of 600 individuals, the number of species collected, as estimated by rarefaction, in the river is 46, compared to 58 species in streams.

**Table 1.** Comparison of the mean insect abundance, species richness and diversity per sample among different habitats and river mesohabitats. Mean values and ranges (in parentheses) and tests of significance are given. Generalized linear models were used for test at the level of habitats and generalized linear mixed models at the level of mesohabitat to account for the effect of site as a random factor. The Simpson’s diversity index is expressed as the inverse of the Simpson’s dominance index. For the comparison at the level of habitats, river samples from a fast section and an adjacent slow section were averaged for each site (wood samples excluded) to provide values comparable to the streams and pools where no mesohabitat distinction was made during sampling.

| Dependent variable | Mean value (range) |  |  | Test of significance |  |
| --- | --- | --- | --- | --- | --- |
|  | Habitat |  |  |  |  |
| | River | Streams | Pools | $F_{2,13}$ | P |
| Mean abundance per sample | 162.7 (125-205) | 91.0 (62-128) | 83.0 (77-89) | 11.89 | **0.002** |
| Mean species richness per sample | 32.0 (29-41) | 22.2 (18-26) | 19.0 (16-22) | 12.62 | **0.001** |
| Diversity (Simpson) | 7.33 (5.84-8.15) | 10.22 (6.93-13.21) | 7.68 (5.41-9.95) | 4.32 | **0.041** |
| Diversity (Shannon) | 2.47 (2.28-2.59) | 2.58 (2.34-2.75) | 2.36 (2.29-2.43) | 2.64 | 0.116 |
| Equitability (Shannon) | 0.72 (0.66-0.75) | 0.84 (0.80-0.88) | 0.81 (0.74-0.88) | 11.17 | **0.002** |
|  | River mesohabitat |  |  |  |  |
| | Fast | Slow | Wood | $\chi^2_2$ | P |
| Total insect abundance | 189.2 (155-247) | 136.2 (84-237) | 128.2 (52-193) | 70.34 | **<0.001** |
| Total insect species richness | 13.4 (11-15) | 18.6 (15-28) | 11.0 (7-13) | 10.33 | **0.006** |
| Diversity (Simpson) | 4.75 (3.05-8.20) | 6.62 (1.87-11.10) | 2.34 (1.55-4.60) | 7.25 | **0.027** |
| Diversity (Shannon) | 1.80 (1.44-2.29) | 2.1 (1.21-2.69) | 1.20 (0.83-1.08) | 9.58 | **0.008** |
| Equitability (Shannon) | 0.69 (0.56-0.85) | 0.72 (0.45-0.88) | 0.51 (0.41-0.75) | 6.87 | **0.032** |

**Fig. 2.**
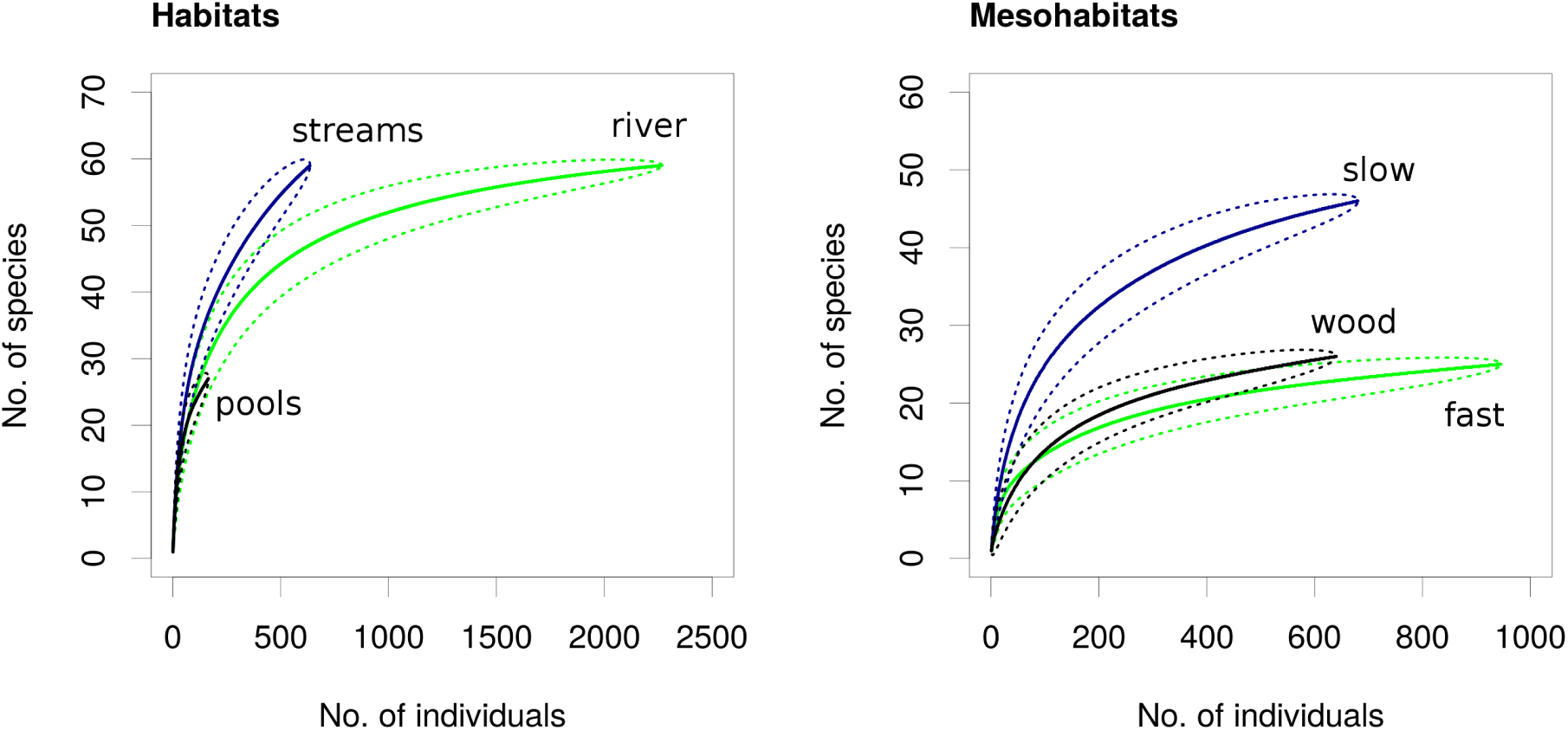
Individual-based rarefaction curves for individual habitats and mesohabitats. The estimated numbers of morphospecies (solid black lines) with 95% confidence limits (dotted black lines) are plotted.

In the river, mean insect abundance per sample was highest in the fast sections (189 individuals, range from 155 to 247) but the slow sections had the highest morphospecies richness per sample (19 morphospecies, range from 15 to 28) (Table 1). Moreover, the slow sections can be supposed to have more species than we collected because the rarefaction curve apparently did not reach the asymptote (Fig. 2). Diversity measured by different indices gave slightly inconsistent results (Table 1) because different indices differently incorporate information on species richness and relative abundance of species in the assessment of diversity. The streams were the most diverse habitat according to Simpson’s index, while Shannon’s diversity index yielded an insignificant result. However, Shannon’s equitability index, which corrects for differences in species richness and presents only information on relative abundances, showed clear differences between habitat types. In the river, the slow sections had the highest diversity followed by the fast sections and wood samples; although the differences were modest (Table 1).

### Assemblage structure

The taxonomical composition of aquatic insect assemblages, in terms of relative abundance of individual morphospecies, differed significantly among the three sampled habitats (RDA, F=2.69, P<0.001, 32.5% explained variance, Fig. 3) as well as among the three mesohabitats in the river (RDA, F=7.65, P<0.001, 65.7% explained variance, Fig. 3). Differences in habitat preferences among major predatory orders – Coleoptera, Odonata and Hemiptera were detected. Dytiscid beetles were equally abundant in all habitats (Supplementary Table S2) but made up the highest proportion of insects in the pools, while Hemiptera and larvae of Odonata dominated in the streams and in the river. Bugs from the families Naucoridae and Aphelocheiridae had higher relative abundance in the fast sections of the river, while larvae of Odonata were more represented in the slow sections. Only two species of dytiscid beetles were found in the river and clearly preferred the slow sections. Beetles from the family Elmidae were almost completely restricted to submerged wood, where they were accompanied by larvae of psephenid beetles and a few species of mayflies and damselflies (Fig. 3, Supplementary Table S2). Ephemeroptera were most represented in the river, where one morphospecies reached higher relative abundance in the slow sections and a second species was more represented in the fast sections and on the submerged wood (Fig. 3, Supplementary Table S2).

**Fig. 3.**
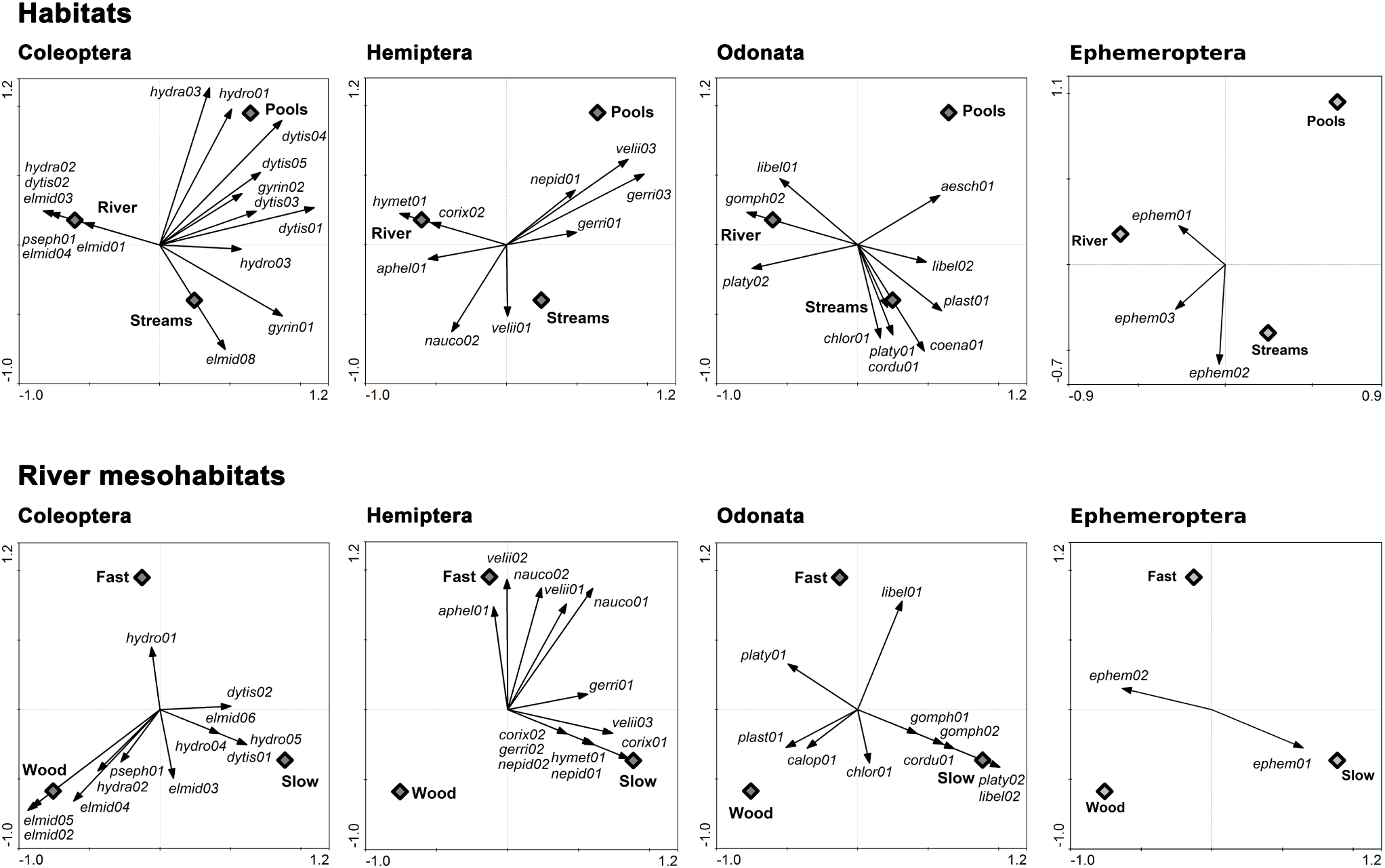
Ordination diagrams displaying the differences in the assemblage composition among different habitats and river mesohabitats (RDA; 1^st^ and 2^nd^ axis). Two ordination analyses were performed – one at the level of habitats and the other at the level of mesohabitats; all insect species were included in both cases. Major groups of aquatic insects are displayed separately only to facilitate visual comparison (species with low fit omitted). Morphospecies codes are explained in Supplementary Table S1. Axes display the ordination scores of individual species, shown by arrows. The relationship between relative species abundance and the habitat types is captured by the directions of the arrows.

The relative abundance of functional feeding groups varied significantly among different habitats (RDA, F=2.58, P=0.033, 31.9% explained variance) and among river mesohabitats (RDA, F=12.30, P<0.001, 75.5% explained variance) (Table 2). Except wood samples, which were overwhelmingly dominated by xylobiont beetles (73.5% of individuals), all habitats were dominated by predators (river – 71.2%, streams – 65.0%, pools – 70.2%), followed by scrapers in running waters (river – 15.6%, streams – 19.1%) and collectors in pools (23.6%). In the river, predators made up larger proportion of the assemblage of the fast compared to slow sections (76.6% vs. 56.3%); scrapers exhibited the opposite pattern (8.7% vs. 29.9%). Shredders were rare reaching the relative abundance only 0.6-11.1% with maximum in the fast sections of the river (Table 2). Assemblage dissimilarity, measured as 1 - the Chao-Sorensen index of similarity, seemed to be highest in the streams and in the slow sections of the river. There was also much higher spread of dissimilarity values in the streams and slow sections of the river, so some sample pairs had almost identical morphospecies composition and some were very different, unlike in the remaining (meso)habitats (Fig. 4).

**Table 2.**
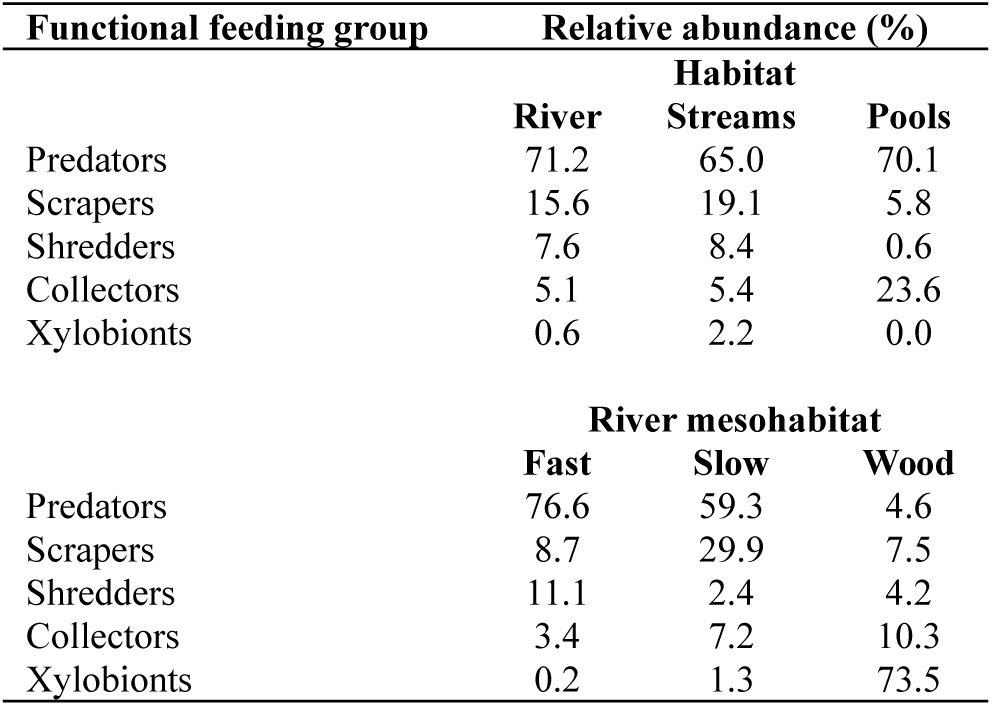
The relative abundance of functional feeding groups in individual habitats and river mesohabitats. The classification of species into functional feeding groups is based on crude literature data because Papua New Guinean freshwater invertebrates are generally poorly known, so caution is necessary when interpreting the numbers.

| Functional feeding group | Relative abundance (%) |  |  |
| --- | --- | --- | --- |
|  | Habitat |  |  |
|  | River | Streams | Pools |
| Predators | 71.2 | 65.0 | 70.1 |
| Scrapers | 15.6 | 19.1 | 5.8 |
| Shredders | 7.6 | 8.4 | 0.6 |
| Collectors | 5.1 | 5.4 | 23.6 |
| Xylobionts | 0.6 | 2.2 | 0.0 |
| River mesohabitat |  |  |  |
|  | Fast | Slow | Wood |
| Predators | 76.6 | 59.3 | 4.6 |
| Scrapers | 8.7 | 29.9 | 7.5 |
| Shredders | 11.1 | 2.4 | 4.2 |
| Collectors | 3.4 | 7.2 | 10.3 |
| Xylobionts | 0.2 | 1.3 | 73.5 |

**Fig. 4.**
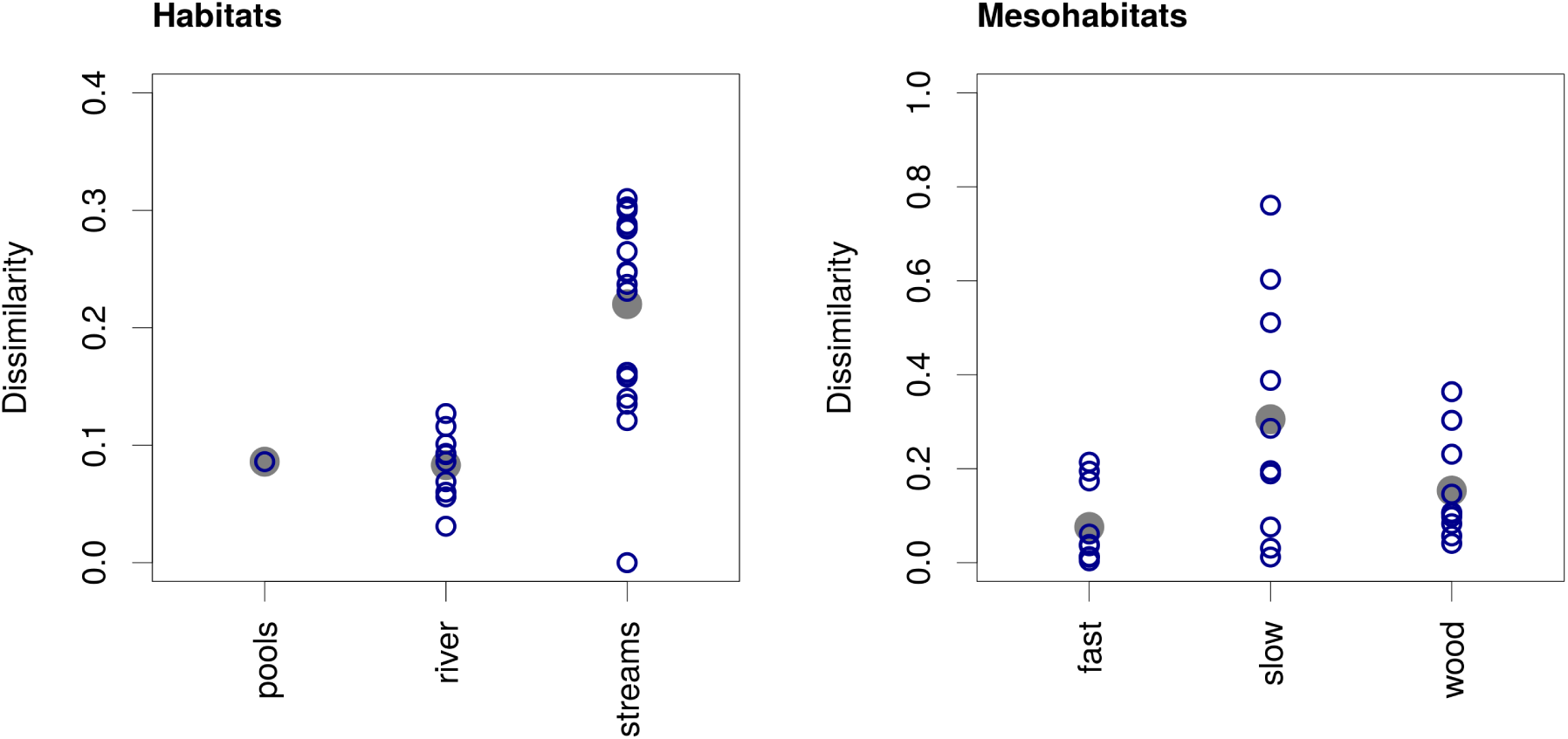
Assemblage dissimilarity in different habitats (a) and mesohabitats (b). Dissimilarity values calculated as1 - Chao-Sorensen index of similarity for all sample pairs within individual habitats are plotted (small empty circles) together with mean values (large grey circles).

## Discussion

This is apparently only the second study on the ecology of aquatic insect assemblages of the island of New Guinea. In his pioneer survey, Dudgeon (1990, 1994) sampled six streams and rivers in an agricultural landscape within the catchments of the Sepik and Ramu Rivers. Our study site is located in the catchment of the Ramu River, but within an area of almost intact lowland rainforest.

### Richness and diversity

The total number of morphospecies collected is comparable to similar studies but the assemblage composition is unusual. Larvae of Diptera, Ephemeroptera, Plecoptera and Trichoptera dominate invertebrate assemblages of most running waters all over the world. Accordingly, Dudgeon (1994) found benthic assemblages of six Papua New Guinean lowland streams (in total 64 morphospecies) to be dominated by Ephemeroptera. However, the Wanang River and nearby streams and pools were clearly dominated by Coleoptera, Hemiptera and Odonata (Supplementary Table S2). Some species of beetles could be counted twice – as adults and larvae. However, only eight morphospecies of beetle larvae were collected (Supplementary Table S2), and it was clear that most of them did not belong to any adults collected. Morphospecies richness of beetles thus could be overestimated at most by a few species, which would have negligible effect on the results. In general, it is possible that the use of morphospecies led to an underestimation of total species richness of the habitats. However, the relative comparison between different habitats and mesohabitats should remain valid.

The comparison of the rarefaction curves clearly showed that the number of morphospecies collected in the river and pools was close to the asymptote but considerably more morphospecies could be found in the streams, which would turn them into the richest habitat (Fig. 2). The likely cause is higher diversity of environmental conditions among different streams than among different sites in the river. In the river, the slow sections had higher mean insect abundance per sample as well as morphospecies richness compared to the fast sections. The pattern is usually reversed in temperate rivers, where riffles harbour higher density, biomass and species richness than slow sections, which is usually ascribed to higher physical habitat heterogeneity providing refuges and higher availability of food in riffles (e.g. Slobodchikoff & Parrot 1977; Brown & Brussock 1991; Lemly & Hilderbrand 2000). This is often the case of mountain rivers, where sediments deposited in slow sections are frequently swept away during high discharge, which precludes the development of sediment-associated fauna (Brown & Brussock 1991). Higher abundance and species richness in slow sections was reported from lowland temperate rivers with higher quantities of stable soft sediments suitable for many benthic invertebrates (McCulloch 1986). Comparable data from tropical streams are rare but most studies found higher abundance and species richness in riffles (e.g. Yule 1996b; Cheshire et al. 2005; Jung et al. 2008). The slow sections of the Wanang River could harbour more insects than the riffles because they accumulated considerable amounts of leaf litter and decaying wood, which could provide rich food base boosting the richness of the whole assemblage (Angradi 1996; Lemly & Hilderbrand 2000).

As expected, submerged wood provided specific environment with low number of specialised morphospecies, which were never or only rarely found in the other mesohabitats. Submerged wood is rarely sampled in studies of running waters and is viewed mostly as a modifier of stream morphology which affects the structure of invertebrate assemblages indirectly by changing flow pattern and by creating debris-dam pools (Lemly & Hilderbrand 2000). By modifying habitat structure and creating accumulations of detritus, large pieces of wood can affect macroinvertebrate diversity and community structure with implications for ecosystem functioning, such as altering rates of accumulation and breakdown of leaf litter (Flores et al. 2011). The lack of knowledge of wood-associated insect fauna hinders any comparison, but e.g. in central Europe, there are only a few rare species specialised on living on submerged wood (Moog 2002). However, Johnson et al. (2003) found nine invertebrate taxa dominantly associated with wood, six of them unique to wood samples, in streams in Minnesota and 35 wood associated species, 27 of them unique to wood, in streams in Michigan. It demonstrates that wood may be an important habitat for aquatic invertebrates, which is corroborated by our results from the Wanang River. More attention should be paid to submerged wood and other neglected habitats such as shore rootlets (Wood & Sites 2002) because they may host distinct assemblages of invertebrates that are overlooked when only bottom fauna is sampled.

### Assemblage structure

Shredders and collectors (filtrators and gatherers) should dominate communities of headwater streams and collectors and scrapers should prevail in medium-sized rivers according to the traditional River Continuum Concept (Vannote et al. 1980). This concept, however, stems from the research of temperate, usually mountainous streams (Vannote et al. 1980). The pattern of assemblage structure in different habitats in the catchment of the Wanang River significantly deviates from these predictions.

The abundance and ecological significance of shredders in tropical compared to temperate streams is a matter of ongoing debate (Boyero et al. 2009). Shredders have been repeatedly shown to be very scarce in tropical streams (Dudgeon 1994; Yule 1996c; Boyero et al. 2009; Li & Dudgeon 2009). Dudgeon (1994) reported relative abundance of shredders of only 0.4% from six Papua New Guinean streams and Yule (1996c) classified only 1.7% of insects in a headwater stream in Bougainville Island as shredders. Shredders were extremely rare also in all habitats we sampled in the catchment of the Wanang River. Toughness and unpalatability of leaves of tropical trees is usually suspected to prevent the invertebrate shredders to thrive in tropical waters and their role in litter breakdown may be taken over by microorganisms (Graça 2001; Boyero et al. 2009). However, recent research showed that paucity of shredders in tropical streams could be partly caused by incorrect classification of species to functional feeding groups (Cheshire et al. 2005). Furthermore, different groups of invertebrates (e.g. snails, crabs and semiaquatic cockroaches) can take up the role of shredders in tropical waters, where shredders typical for temperate streams, e.g. stoneflies and caddisflies, are rare (Yule et al. 2009). More detailed research including thorough examination of feeding habits of individual species (Cheshire et al. 2005, Li & Dudgeon 2009) will be needed to evaluate the role of different organisms for leaf litter breakdown in Papua New Guinean waters.

The high proportion of predators we detected in the catchment of the Wanang River (overall 57% abundance) is unprecedented. Dudgeon (1994) reported only 7% of individuals belonging to predators in his lowland Papua New Guinean streams. Cheshire et al. (2005), who reported predators making up about 25% of insect abundance in two tropical streams in Queensland, speculated that their populations could be sustained by high productivity of their prey. However, intraguild predation and cannibalism are common in aquatic insects including dytiscid beetles (Yee 2010) and odonate larvae (McPeek 1990; Johansson 1991, 1993) and could provide an important food source for maintaining high predator densities.

Predatory taxa also exhibited different distribution among the habitats and mesohabitats (Fig. 3). These differences may reflect contrasting habitat or prey preferences (Klecka & Boukal 2012) or avoidance of interference and intraguild predation. These mechanisms are known to drive habitat partitioning among competing predatory caddisflies and stoneflies in temperate streams (Sircom & Walde 2009) and among backswimmers (*Notonecta*) in stagnant waters (Giller & McNeill 1981). Furthermore, dytiscid beetles may be rare in waters dominated by dragonflies as a consequence of intraguild predation (Larsson 1990). Contrasting habitat preferences could also result from the effects of water availability on insects with different life histories (Stoks & McPeek 2003). Dytiscidae and Heteroptera have rapid larval development and can utilise temporary habitats. On the other hand, Odonata have often long larval period and therefore should prefer more stable habitats. Correspondingly, Dytiscidae were equally abundant in the river, streams and pools (Supplementary Table S2) and made up the highest proportion of the insect assemblage in the pools, which are prone to drying out, whereas Odonata (but also Heteroptera) were less represented in the pools (Fig. 3). Several of these factors could act together and create a complex environmental gradient (Stoks and McPeek 2003).

Some aspects of the community composition could be affected by the methods used in the field sampling. Specifically, we used a relatively coarse mesh size (0.75 mm), while the mesh size of more often used sampling devices such as a handnet and a Surber sampler is usually 0.3 – 0.5 mm. Consequently, very small species of aquatic insects, such as young larvae of Diptera, could have been missed during our sampling. However, it is unlikely that the high abundance of predatory Heteroptera, Coleoptera and Odonata in the Wanang River could be attributed simply to sampling bias. Moreover, we could have missed some large mobile species able to avoid capture by rapid escape (Klecka & Boukal 2011), which are usually predatory, making overestimation of the importance of predators in this study unlikely. In addition, Barba et al. (2010) found that although mesh size can affect some metrics of invertebrate community structure, species richness and differences among sampling sites are robust to mesh size. On a more detailed level, we could have underestimated the diversity of Diptera, which contain species with very small aquatic larvae. Contrary to the sampling methods, analysis of assemblage structure is robust to the use of morphospecies. Even though true species richness might be underestimated, functional groups would be unaffected because they are defined at a coarser taxonomical level.

### Conclusions

Overall, the results show intriguing departures from common patterns of the structure of aquatic insect assemblages known from more thoroughly studied regions. Freshwater habitats I studied in a Papua New Guinean lowland rainforest harboured relatively rich insect assemblages. The aquatic insects exhibited distinct habitat and mesohabitat selectivity and their assemblages were dominated by predators (beetles, bugs and dragonflies), which is very unusual. Terrestrial ecologists have been studying rich tropical forests of Papua New Guinea for a long time (e.g. Novotny et al. 2002). Its understudied freshwaters surely deserve similar attention; especially in the light of ongoing debates on characteristic properties of tropical streams and their similarities to and differences from temperate streams (e.g. Boyero et al. 2009).

## Acknowledgements

I thank Robin Kalwa, Maling Rimandai and Manuel Simbai for field assistance. Vojtech Novotny, Jan Leps and the staff of New Guinea Binatang Research Center (Madang, Papua New Guinea) provided important logistical support. I would like to thank the reviewers for valuable comments on the manuscript. This research was performed during a field course of tropical ecology organized and partly financed by the Faculty of Science, University of South Bohemia (Ceske Budejovice, Czech Republic), Binatang Research Center (Madang, Papua New Guinea) and PNG Institute of Biological Research (Goroka, Papua New Guinea). This research was conducted with institutional support RVO:60077344.

## Supplementary data

Aquatic insects of a lowland rainforest in Papua New Guinea: assemblage structure in relation to habitat type

Jan Klecka

Laboratory of Theoretical Ecology, Biology Centre of the Academy of Sciences of the Czech Republic v.v.i., Institute of Entomology, Branišovská 31, České Budějovice, Czech Republic,

**Table S1.** List of sampling sites. Site no. refers to Fig. 1. Mean width is given for the river and streams and estimated area for the stagnant pools. The bottom substrate was classified as soft sediment (particle size<0.5mm), sand (0.5–5 mm), gravel (5–50 mm) and cobbles (50–200 mm); leaves refer to a layer of decaying tree leaves. Exposure to sunshine was classified based on relative site area shaded at noon.

| Site no. | Habitat | Mesohabitat | Width (m)/area (m<sup>2</sup>) | Depth (m) | Bottom substrate | Exposure |
| --- | --- | --- | --- | --- | --- | --- |
| 1 | pools |  | 1.0, 2.5 | 0.6 | soft sediment, sand, leaves | partly shaded |
| 2 | stream |  | 1.0 | 0.5 | sand, gravel, leaves | shaded |
| 3 | stream |  | 0.5 | 0.3 | gravel | shaded |
| 4 | stream |  | 0.5 | 0.2 | gravel, cobbles, leaves | shaded |
| 5 | river | fast | 7.0 | 0.2 | cobbles | exposed |
|  |  | slow, wood | 7.0 | 1.0 | soft sediment, leaves | shaded |
| 6 | river | fast | 8.0 | 0.1 | gravel | partly shaded |
|  |  | slow, wood | 5.0 | 1.0 | soft sediment, sand, leaves | partly shaded |
| 7 | river | fast | 5.0 | 0.2 | cobbles | partly shaded |
|  |  | slow, wood | 6.0 | 0.5 | soft sediment, sand, gravel, leaves | shaded |
| 8 | stream |  | 0.5 | 0.5 | gravel, cobbles | partly shaded |
| 9 | stream |  | 1.0 | 0.2 | gravel, cobbles, leaves | exposed |
| 10 | stream |  | 2.5 | 0.5 | gravel, cobbles, leaves | shaded |
| 11 | pools |  | 1.0, 1.5, 1.5, 2.0 | 0.2 | gravel, leaves | shaded |
| 12 | river | fast | 6.0 | 0.1 | gravel, cobbles | partly shaded |
|  |  | slow, wood | 6.0 | 0.3 | gravel | exposed |
| 13 | river | fast | 5.0 | 0.1 | gravel, cobbles | exposed |
|  |  | slow, wood | 4.0 | 1.0 | soft sediment, leaves | exposed |
| 14 | stream |  | 2.0 | 0.3 | sand, gravel, leaves | partly shaded |

**Table S2.** List of morphospecies. The total numbers of specimens collected in individual habitats and mesohabitats are given for all morphospecies. Taxa are listed alphabetically within each taxonomical level. In Coleoptera, A and L in parentheses denote the stage collected (adult, larva). *N* = total number of individuals collected.

| Order and family | Species code | N | Habitats |  |  | River mesohabitats |  |  |
| --- | --- | --- | --- | --- | --- | --- | --- | --- |
|  |  |  | River | Streams | Pools | Fast | Slow | Wood |
| Coleoptera |  |  |  |  |  |  |  |  |
| Dytiscidae (A) | dytis01 | 76 | 2 | 35 | 39 | 0 | 2 | 0 |
| Dytiscidae (A) | dytis02 | 59 | 59 | 0 | 0 | 4 | 55 | 0 |
| Dytiscidae (A) | dytis03 | 19 | 0 | 10 | 9 | 0 | 0 | 0 |
| Dytiscidae (A) | dytis04 | 7 | 0 | 2 | 5 | 0 | 0 | 0 |
| Dytiscidae (A) | dytis05 | 11 | 0 | 6 | 5 | 0 | 0 | 0 |
| Elmidae (A) | elmid01 | 2 | 2 | 0 | 0 | 2 | 0 | 0 |
| Elmidae (A+L) | elmid02 | 452 | 451 | 1 | 0 | 0 | 0 | 451 |
| Elmidae (L) | elmid03 | 6 | 6 | 0 | 0 | 0 | 3 | 3 |
| Elmidae (A) | elmid04 | 15 | 15 | 0 | 0 | 0 | 2 | 13 |
| Elmidae (L) | elmid05 | 28 | 28 | 0 | 0 | 0 | 0 | 28 |
| Elmidae (A) | elmid06 | 3 | 2 | 1 | 0 | 0 | 2 | 0 |
| Elmidae (L) | elmid07 | 1 | 0 | 1 | 0 | 0 | 0 | 0 |
| Elmidae (L) | elmid08 | 11 | 0 | 11 | 0 | 0 | 0 | 0 |
| Gyrinidae (A) | gyrin01 | 33 | 0 | 29 | 4 | 0 | 0 | 0 |
| Gyrinidae (A) | gyrin02 | 2 | 0 | 1 | 1 | 0 | 0 | 0 |
| Hydraenidae (A) | hydra01 | 5 | 4 | 1 | 0 | 1 | 3 | 0 |
| Hydraenidae (A) | hydra02 | 17 | 17 | 0 | 0 | 0 | 1 | 16 |
| Hydraenidae (A) | hydra03 | 15 | 8 | 0 | 7 | 7 | 1 | 0 |
| Hydraenidae (A) | hydra04 | 3 | 0 | 3 | 0 | 0 | 0 | 0 |
| Hydrophilidae (L) | hydro01 | 14 | 5 | 1 | 8 | 5 | 0 | 0 |
| Hydrophilidae (A) | hydro02 | 5 | 2 | 3 | 0 | 0 | 1 | 1 |
| Hydrophilidae (A) | hydro03 | 4 | 0 | 3 | 1 | 0 | 0 | 0 |
| Hydrophilidae (A) | hydro04 | 31 | 27 | 2 | 2 | 0 | 27 | 0 |
| Hydrophilidae (A) | hydro05 | 3 | 1 | 2 | 0 | 0 | 1 | 0 |
| Hydrophilidae (A) | hydro06 | 1 | 0 | 1 | 0 | 0 | 0 | 0 |
| Psephenidae (L) | pseph01 | 20 | 20 | 0 | 0 | 0 | 3 | 17 |
| Scirtidae (L) | scirt01 | 3 | 2 | 1 | 0 | 0 | 1 | 1 |
| Diptera |  |  |  |  |  |  |  |  |
| Chironomidae | chiro01 | 7 | 7 | 0 | 0 | 0 | 7 | 0 |
| Chironomidae | chiro02 | 11 | 1 | 9 | 1 | 0 | 1 | 0 |
| Culicidae | culic01 | 29 | 10 | 1 | 18 | 0 | 10 | 0 |
| ? | dipt01 | 22 | 20 | 2 | 0 | 14 | 1 | 5 |
| ? | dipt02 | 2 | 0 | 2 | 0 | 0 | 0 | 0 |
| ? | dipt03 | 7 | 5 | 2 | 0 | 3 | 2 | 0 |
| ? | dipt04 | 1 | 0 | 1 | 0 | 0 | 0 | 0 |
| ? | dipt05 | 2 | 1 | 1 | 0 | 1 | 0 | 0 |
| ? | dipt07 | 1 | 0 | 1 | 0 | 0 | 0 | 0 |
| ? | dipt08 | 1 | 0 | 1 | 0 | 0 | 0 | 0 |
| ? | dipt09 | 1 | 1 | 0 | 0 | 0 | 0 | 1 |
|  |  |  | River | Streams | Pools | Fast | Slow | Wood |
| Ephemeroptera |  |  |  |  |  |  |  |  |
| Caenidae | caeni01 | 156 | 134 | 14 | 8 | 5 | 125 | 4 |
| Caenidae | caeni02 | 218 | 126 | 90 | 2 | 66 | 16 | 44 |
| Caenidae | caeni03 | 37 | 25 | 12 | 0 | 11 | 12 | 2 |
| Prosopistomatidae | proso01 | 1 | 1 | 0 | 0 | 0 | 1 | 0 |
| Hemiptera |  |  |  |  |  |  |  |  |
| Aphelocheiridae | aphel01 | 18 | 15 | 3 | 0 | 15 | 0 | 0 |
| Corixidae | corix01 | 51 | 51 | 0 | 0 | 0 | 51 | 0 |
| Corixidae | corix02 | 31 | 31 | 0 | 0 | 0 | 31 | 0 |
| Gerridae | gerri01 | 34 | 13 | 16 | 5 | 4 | 9 | 0 |
| Gerridae | gerri02 | 5 | 3 | 2 | 0 | 0 | 3 | 0 |
| Gerridae | gerri03 | 12 | 0 | 6 | 6 | 0 | 0 | 0 |
| Hydrometridae | hymet01 | 3 | 3 | 0 | 0 | 0 | 3 | 0 |
| Mesoveliidae | mesov01 | 6 | 4 | 2 | 0 | 0 | 3 | 1 |
| Naucoridae | nauco01 | 620 | 543 | 75 | 2 | 378 | 156 | 9 |
| Naucoridae | nauco02 | 103 | 71 | 31 | 1 | 55 | 16 | 0 |
| Nepidae | nepid01 | 8 | 2 | 3 | 3 | 0 | 2 | 0 |
| Nepidae | nepid02 | 5 | 3 | 2 | 0 | 0 | 3 | 0 |
| Notonectidae | noton01 | 1 | 0 | 1 | 0 | 0 | 0 | 0 |
| Veliidae | velii01 | 175 | 100 | 67 | 8 | 71 | 28 | 1 |
| Veliidae | velii02 | 154 | 150 | 4 | 0 | 143 | 7 | 0 |
| Veliidae | velii03 | 41 | 10 | 16 | 15 | 1 | 9 | 0 |
| Megaloptera |  |  |  |  |  |  |  |  |
| ? | megal01 | 1 | 1 | 0 | 0 | 0 | 0 | 1 |
| Odonata |  |  |  |  |  |  |  |  |
| Aeschnidae | aesch01 | 2 | 0 | 1 | 1 | 0 | 0 | 0 |
| Calopterygidae | calop01 | 1 | 1 | 0 | 0 | 0 | 0 | 1 |
| Chlorophycidae | chlor01 | 17 | 8 | 9 | 0 | 0 | 4 | 4 |
| Coenagrionidae | coena01 | 19 | 0 | 19 | 0 | 0 | 0 | 0 |
| Corduliidae | cordu01 | 22 | 10 | 11 | 1 | 0 | 10 | 0 |
| Gomphidae | gomph01 | 6 | 5 | 1 | 0 | 0 | 5 | 0 |
| Gomphidae | gomph02 | 4 | 4 | 0 | 0 | 0 | 4 | 0 |
| Libellulidae | libel01 | 78 | 64 | 7 | 7 | 51 | 13 | 0 |
| Libellulidae | libel02 | 45 | 19 | 21 | 5 | 0 | 19 | 0 |
| Platystictidae | plast01 | 18 | 6 | 11 | 1 | 1 | 0 | 5 |
| Platycnemididae | platy01 | 27 | 6 | 21 | 0 | 4 | 0 | 2 |
| Platycnemididae | platy02 | 20 | 16 | 4 | 0 | 0 | 16 | 0 |
| Plecoptera |  |  |  |  |  |  |  |  |
| ? | pleco01 | 1 | 0 | 1 | 0 | 0 | 0 | 0 |
| Trichoptera |  |  |  |  |  |  |  |  |
| ? | trich01 | 130 | 102 | 28 | 0 | 96 | 1 | 5 |
| ? | trich02 | 38 | 15 | 22 | 1 | 1 | 8 | 6 |
| ? | trich03 | 13 | 11 | 2 | 0 | 0 | 1 | 10 |
| ? | trich04 | 10 | 9 | 1 | 0 | 0 | 2 | 7 |
| ? | trich05 | 6 | 6 | 0 | 0 | 6 | 0 | 0 |
| ? | trich06 | 4 | 4 | 0 | 0 | 1 | 0 | 3 |
| Total number of individuals |  | 3071 | 2268 | 637 | 166 | 946 | 681 | 641 |
| Total number of species |  | 78 | 59 | 59 | 27 | 25 | 46 | 26 |

**Fig. S1.**
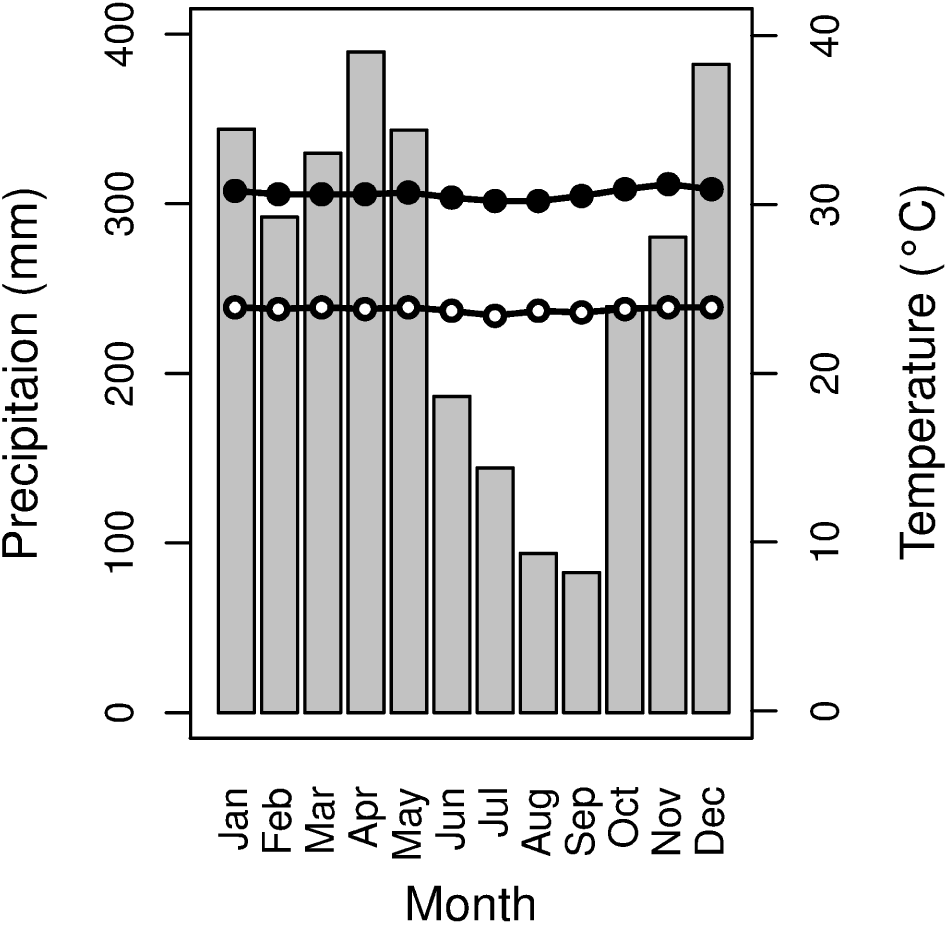
Mean monthly temperature (filled circles = mean daily maximum, empty circles = mean daily minimum) and rainfall (grey vertical bars) for Madang, Papua New Guinea (monthly averages for years 1973–2006). Data were obtained from PNG National Weather Service.

